# Proteolytic activation of fatty acid synthase signals pan-stress resolution

**DOI:** 10.1101/2023.09.27.559743

**Authors:** Hai Wei, Yi M. Weaver, Chendong Yang, Yuan Zhang, Guoli Hu, Courtney M. Karner, Ralph J. DeBerardinis, Benjamin P. Weaver

## Abstract

Chronic stress and inflammation are not only outcomes of pathological states but rather major drivers of many human diseases ^1-4^. Ideally, a given stress program is downregulated to basal levels upon restoration of homeostasis. Chronic responsiveness despite stress mitigation suggests a failure to sense the resolution of the initiating stressor. Here we show that a proteolytic cleavage event of fatty acid synthase (FASN) activates a global cue for stress resolution. FASN is well-established as the multifunctional enzyme catalyzing *de novo* biosynthesis of saturated fatty acid ^5, 6^. Surprisingly, our results demonstrate FASN functioning as a signaling molecule promoting an anti-inflammatory profile apart from fatty acid synthesis. Redox-dependent proteolytic cleavage of FASN by caspase activates a truncated C-terminal enzymatic fragment (FASN-CTF) that is sufficient to down-regulate multiple aspects of stress-responsiveness including gene expression and metabolic programs. Only a fraction of FASN is cleaved allowing for continued fat synthesis. FASN-CTF can signal stress resolution across tissues in a cell non-autonomous manner. Consistent with these findings, FASN processing is also seen in well-fed but not fasted mouse liver. As down-regulation of stress responsiveness is critical to health, our findings provide a potential pathway to control the magnitude for diverse aspects of stress responses.

## Introduction

Alterations in nutrient availability, oxidative imbalances, and exposures to cytotoxic substances or pollutants are all examples of cellular stressors. When faced with these insults, cells mount the relevant stress mitigation program to restore homeostasis. Outcomes include upregulating buffering factors like chaperones, repairing damaged DNA, autophagy of cellular components, or death of irreversibly damaged cells. Whether animals have a globally-acting mechanism to detect the resolution of diverse stressors and signal the down-regulation of multiple features of stress responses is unknown. However, such a mechanism would offer a survival advantage to limit loss of energy stores, speed wound healing and prepare the animal for encounters with another stressor. For multi-cellular organisms, coordinating stress recovery across tissues imposes additional challenges.

Adaptations to stressors not only require altered gene expression but also metabolic rewiring to repurpose energy stores. Metabolic adaptations to stressors can result in the generation of reactive oxygen or nitrogen species (ROS, RNS). These reactive species cause extensive cellular damage including lipid peroxidation resulting in toxic aldehyde intermediates that further damage other macromolecules and lead to death of tissues ^7^. Thus, lipids have an intriguing duality in stress responses as both energy stores and liability to damaging agents resulting in lipotoxicity. This underscores the need to have dynamic control of lipid availability but also safeguard the cell.

Cytosolic fatty acid synthase (FASN) is the biosynthetic enzyme responsible for the *de novo* synthesis of saturated fatty acids, the building blocks of fats and higher-order lipids ^5, 6^. Fat synthesis by FASN is a major pathway consuming energy stores in the form of NADPH. Intriguingly, upregulation of *FASN* expression is a hallmark of tumorigenesis where tumor cells are thought to acquire *de novo* lipid synthesis ^8, 9^. However, it is not clear whether *FASN* upregulation solely reflects increased nutrient demand for growth as lipids are involved in multiple aspects of cellular transformation including immune suppression and drug resistance, thereby preventing death of transformed cells ^9^.

Caspases are well-known for activating pro-inflammatory cytokines or committing cells to programmed cell death if a genotoxic insult is not repaired. This same family of cysteine proteases also has ancient non-lethal roles in promoting differentiation ^10^. However, across metazoans, these same proteases have poorly understood roles in supporting homeostasis where modulating caspase function alters stress resistance in *C. elegans* ^11, 12^, reprograms cardiac hypertrophic responses in rats ^13^, and accelerates aging in mice ^14^. Our previous work identified CED-3 caspase blocking a p38 MAPK-dependent epidermal pathogen response ^12^. Also, the dual-leucine zipper kinase (DLK-1, MAPK3K12) pathway was implicated downstream of caspase in neuronal regeneration following injury ^15^. Additionally, stress-induced caspase-2 cleavage of site 1 protease (S1P) triggers persistent activation of the transcription factor SREBP leading to NASH development ^16^. However, for each of these cases, the caspase was found targeting a factor activating a stress response.

Beyond detecting an insult to activate a stress response, little is known in how animals sense the resolution of stressors. Here we identify a signaling function of FASN that is independent of fatty acid biosynthesis. We show that redox-dependent caspase cleavage of FASN serves as a sensing mechanism to gauge appropriate responses to diverse stressors. With cysteine active sites, the proteolytic activity of caspases can be inhibited by stressors in a redox-dependent manner. When the stressor is mitigated, limited amounts of the C-terminal fragment (FASN-CTF) act as a strong cue to down-regulate widespread features of animal stress responses, analogous to an anti-inflammatory signal. Moreover, FASN-CTF enzymatic activity is required to suppress stress-responsive gene expression programs and promote lipid mobilization. Forced expression of FASN-CTF prior to stress amelioration compromises survival underscoring its function as a stress-resolution cue.

## Results

### Caspase controls magnitude and persistence of diverse stress responses

Disrupting the caspase pathway in *C. elegans* provided us a model to examine chronic stress responses. As early as 4 hours of tunicamycin treatment, the *ced-3(-)* caspase and upstream activating Apaf gene *ced-4(-)* null mutants had dramatically enhanced induction of the Hsp70 family (HSPA5, BiP) ER stress marker HSP-4p::GFP relative to wild-type animals (Fig. 1a,b and Extended Data Fig. 1a,b). This enhanced response persisted out to 2 days (Fig. 1b). Moreover, although the *ced-3(-)* and *ced-4(-)* null mutants normally develop slower than wild type, both mutants had faster development under ER stress conditions compared to wild-type animals (Fig. 1c and Extended Data Fig. 1c) suggesting that the higher induction of HSP-4p::GFP reflects an enhanced survival advantage upon tunicamycin-induced ER stress.

**Fig. 1.**
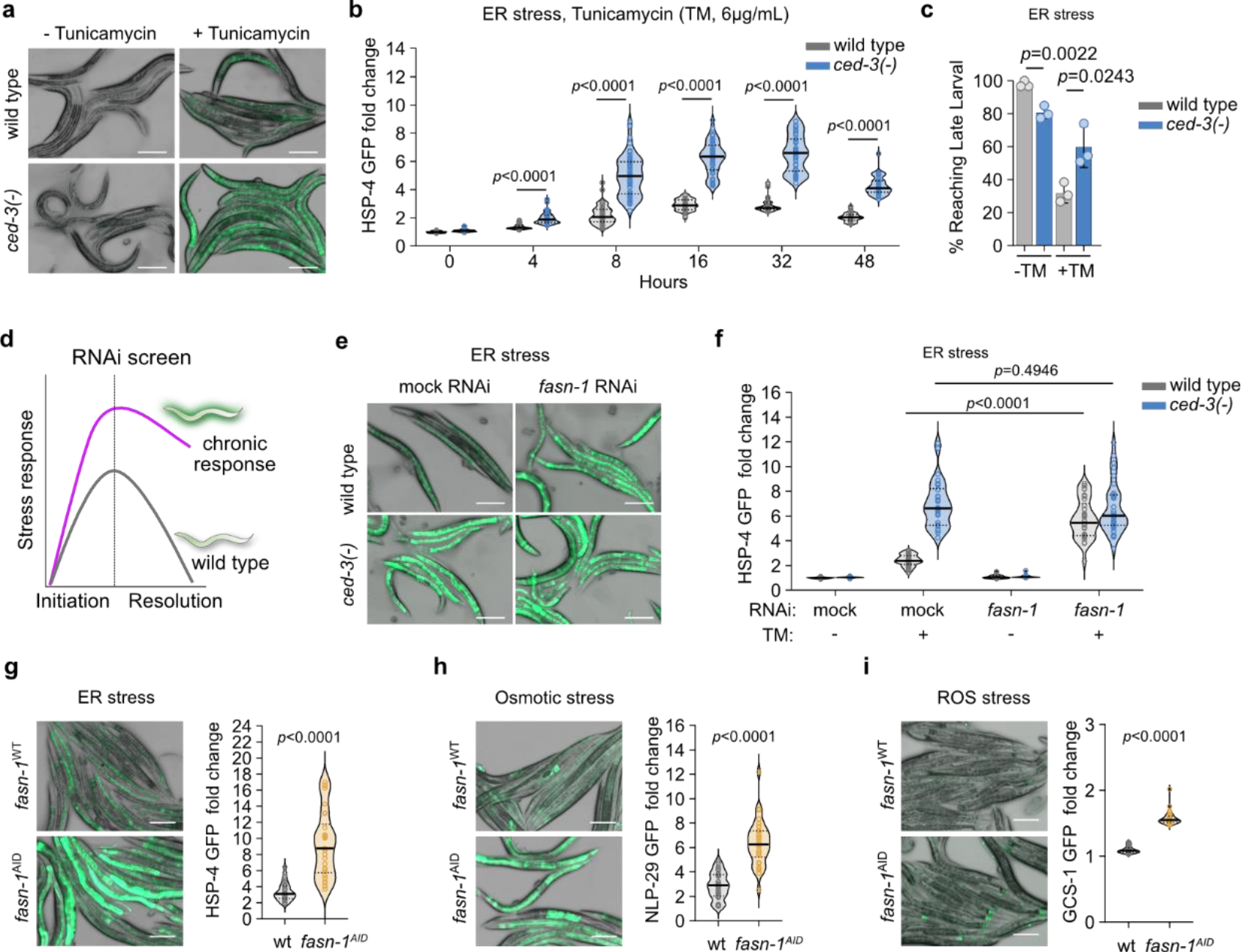
Caspase and Fatty Acid Synthase (FASN) limit chronic pan-stress responses. **a-c**, induction (**a**) and quantification (**b**) of intestinal ER stress reporter HSP-4p::GFP (Hsp70 member 5) with tunicamycin (TM) treatment to induce ER stress. Quantification of animals escaping larval arrest (**c**) with TM treatment. **d-e**, targeted RNAi screen for chronic stress response factors (**d**) reveals *fasn-1* acts like *ced-3* to limit stress response. HSP-4p::GFP induction (**e**) and quantification (**f**). **g-i**, eliminating FASN-1 protein using auxin induced degron (AID) also induces expression of stress reporters under ER (**g**), osmotic (**h**) and ROS (**i**) stresses by. For **a,e,g,h,i**, scale bar, 200μm. For **b,f,g,h,i** each circle represents one animal. Mean pixel intensity of each animal was normalized to mean value of wild type at 0hr (no stress) and plotted as fold change. Violin plots show median (solid line) with quartiles (dashed line). n=30-50 animals for each time point. For **c**, mean±SD, n=3 biological replicates. Total animals for each bar ranges from 141-215, *p* values were calculated using Mann-Whitney test (**b,f,g,h,i**) and unpaired t-test (**c**).

We next sought to test whether *ced-3* and *ced-4* mutants would have similar persistent phenotypes with other stressors. Analogous to the heightened ER stress response, the *ced-3(-)* and *ced-4(-)* null mutants had dramatically enhanced induction of markers for both osmotic stress (NLP29p::GFP) and ROS stress (GCS-1p::GFP) relative to wild-type animals when challenged with high salt and paraquat, respectively (Extended Data Figs. 1d-g and 2a-d).

The *ced-3(-)* and *ced-4(-)* mutants also had enhanced motility when given the salt challenge (Extended Data Fig. 1h,i) and enhanced survival on paraquat relative to wild-type animals (Extended Data Fig. 2e,f). Importantly, for each of the diverse stress challenges, wild-type animals never reached the same pronounced induction as seen with the *ced-3* caspase and *ced-4* Apaf mutants. Also, the *ced-3* and *ced-4* mutants had marked variability for the induction of the stress reporters. Together, these results indicate that caspase pathway normally regulates the overall magnitude of responsiveness to diverse stressors. We were therefore intrigued to identify what target pathway is responsible for the enhanced pan-stress responsiveness

**Fig. 2.**
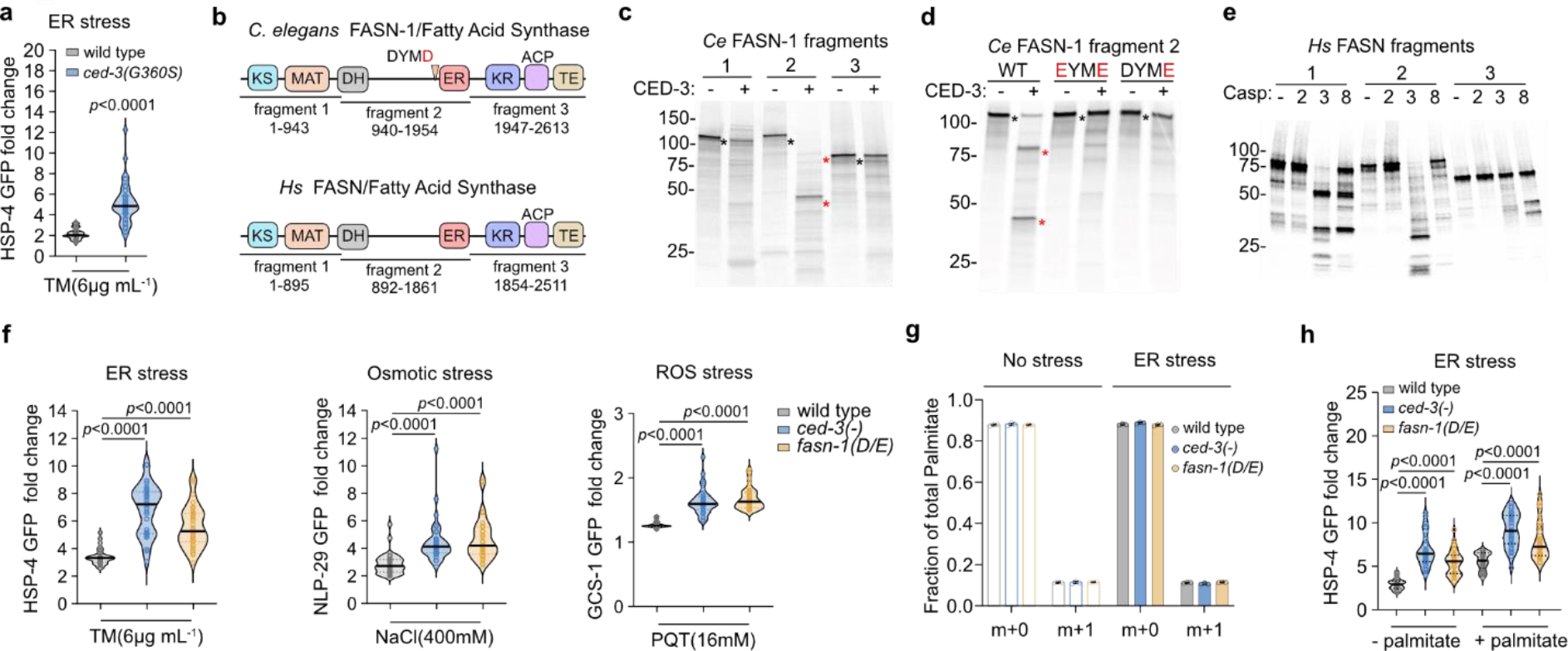
FASN activated by caspase cleavage limits pan-stress responses independent of palmitate synthesis. **a**, induction of HSP-4p::GFP by caspase active-site mutation G360S. **b**, highly conserved FASN domains and fragments used for *in vitro* caspase cleavage analyses. Enzymatic domains indicated with square boxes. **c-d**, CED-3 *in vitro* cleavage of ^35^S-labeled *C. elegans* FASN-1 fragments (**c**), the DYMD to DYME mutation (D/E) in fragment 2 blocks CED-3 cleavage (**d**). Red *, cleaved product. Black *, full-length molecule. **e**, Conserved cleavage. ^35^S-labeled human FASN is also cleaved in fragment 2 by caspase-3 producing a stable C-terminal fragment 3 similar to *C. elegans*. Casp: caspases 2,3,8 as indicated for each lane. **f**, cleavage-resistant *fasn-1(D/E) C. elegans* mutant phenocopies *ced-3(-)* mutant for enhanced inductions of ER, Osmotic and ROS stress reporters. **g**, measurement of palmitate synthesis using 50% deuterated water (D_2_O) labeling for 16 hours in the presence or absence of ER stress. m+0, unlabeled palmitate. m+1, deuterium-labeled palmitate representing newly synthesized palmitate. n=3 biological samples. Mean ± SD. **h**, HSP-4p::GFP induction under ER stress with palmitate supplementation. For **a,f,h**, each circle represents one animal. Mean pixel intensity of each animal was normalized to mean value of wild type without stress and plotted as fold change. Violin plots show median (solid line) with quartiles (dashed line). n=30 animals for each condition. *p* values were calculated using Mann-Whitney test.

### Fatty Acid Synthase (FASN) acts like caspase to limit stress response magnitude

To reveal the caspase signaling target, we performed an RNAi screen using the HSP-4p::GFP reporter with tunicamycin-induced ER stress to identify candidate factors contributing to persistently-heightened stress responses (Fig. 1d). The targeted screen examined established stress-responsive regulators and genes upregulated in stress (Extended Data Fig. 3a-c). The *C. elegans* fatty acid synthase (FASN) ortholog *fasn-1* gene, acted like *ced-3(-)* and *ced-4(-)* mutants with enhanced induction and pronounced variability of HSP-4p::GFP expression under ER stress (Fig. 1e,f and Extended Data Fig. 3c). Moreover, because *fasn-1* RNAi knockdown did not enhance stress response in *ced-3(-)* mutants (Fig. 1f), we wanted to confirm that loss of FASN-1 expression acts like *ced-3(-)* null.

**Fig. 3.**
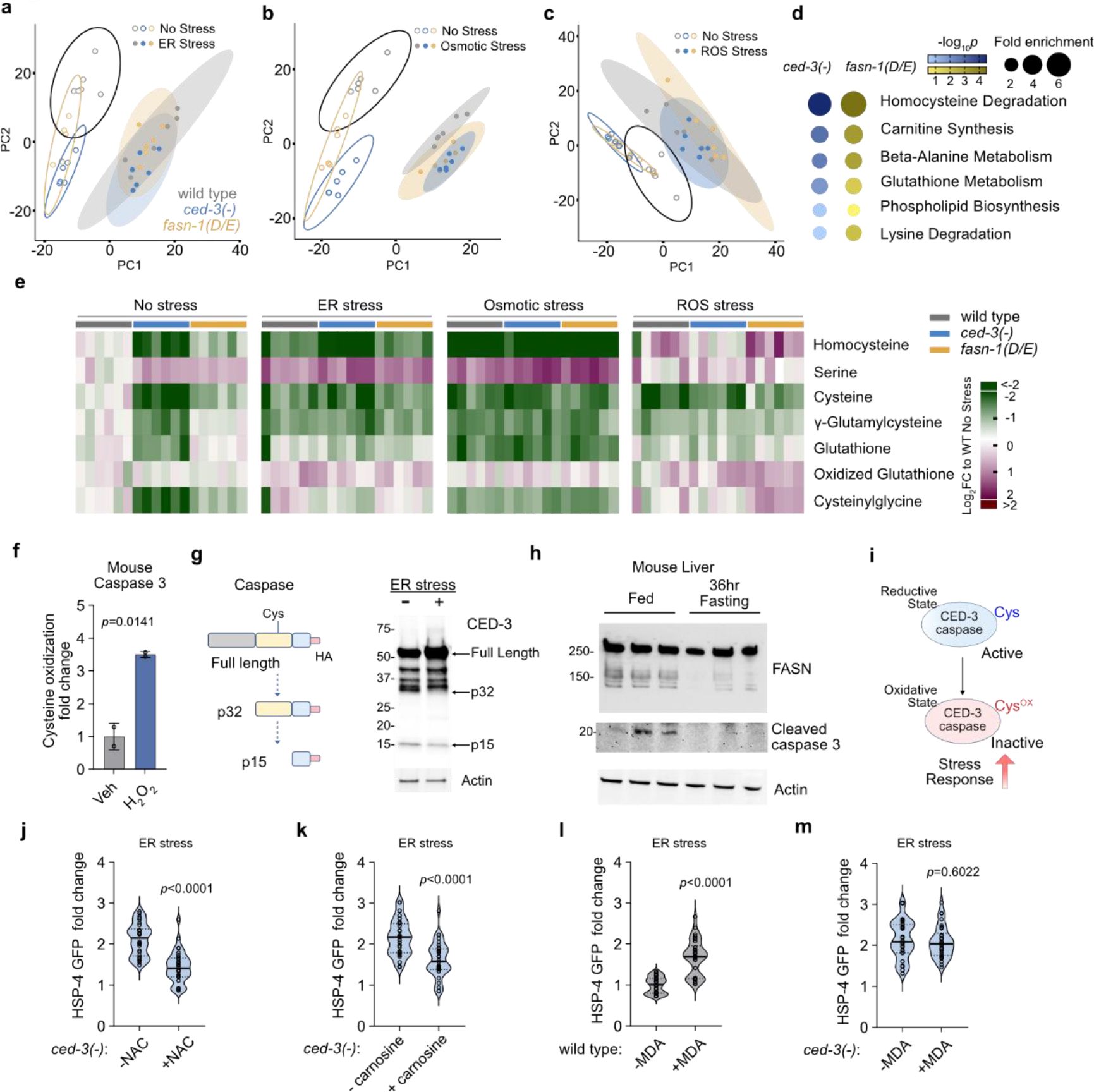
Interplay of cellular redox state and FASN cleavage determines stress responsiveness. **a-c**, principle components analysis (PCA) of untargeted metabolites for wild type, *ced-3(-)* and *fasn-1(D/E)* animals with ER (**a**), osmotic (**b**) and ROS (**c**) stressors, n=6 biological replicates. **d**, metabolite enrichment analyses for *ced-3(-)* null and *fasn-1(D/E)* mutants compared to wild-type using MetaboAnalyst (https://www.metaboanalyst.ca/) with SMPDB. FDR<0.1. **e**, heatmap of detected metabolites in glutathione pathway. Log_2_-fold change relative to mean value of wild type no stress. n=6 biological replicates. **f**, quantification of caspase-3 cysteine oxidation in primary mouse calvarial osteoblasts (cOBs) under H_2_O_2_ treatment using DCP-BIO IP Mass Spec. DCP-Bio1 selectively reacts with cysteine sulfenic acid detecting oxidation products. Fraction of caspase-3 with oxidized cysteine to total caspase-3 was calculated using abundance of DCP-Bio1 labeled protein normalized to abundance of total caspase-3 protein. Veh, water. n=2 biological replicates. **g**, diagram of CED-3 auto-processing monitored by C-term HA tag and western blot of endogenous CED-3 with C-term HA tag revealed decreased caspase auto-processing *in vivo* under ER stress. **h**, Western blot of liver extracts from well-fed or 36 hour fasted mice showing processing of FASN and cleaved caspase-3. n= 3 mice for each condition. **i**, diagram of caspase cys active site as a function of oxidative state and impact on stress responsiveness. **j-k**, Induction of HSP-4p::GFP under ER stress with N-acetylcysteine (NAC) (**j**) or carnosine (**k**) anti-oxidant supplementation in *ced-3(-)* animals. **l,m**, Induction of HSP-4p::GFP under ER stress with oxidized lipid metabolite MDA (malondialdehyde) supplementation in wild type (**l**) and *ced-3(-)* animals (**m**). Each circle represents one animal. Mean pixel intensity of each animal was normalized to mean value of wild type without stress and plotted as fold change. Violin plots show median (solid line) with quartiles (dashed line). n=30-40 animals for each time point. *p* values were calculated using unpaired ttest (**f**) or Mann-Whitney test (**j-m**).

To confirm that loss of FASN-1 function resulted in enhanced stress response in animals with intact CED-3 function, we used an alternative technique to deplete FASN at the protein level. We added HA-labelled auxin-induced degron (AID) tags to both the N- and C-termini of the *fasn-1* coding sequence in the endogenous genomic locus using CRISPR mutagenesis. When treated with auxin, the amount of FASN protein was significantly reduced (Extended Data Fig. 3d) and depletion enhanced induction of HSP-4p::GFP with ER stress (Fig. 1g), NLP-29p::GFP for osmotic stress (Fig. 1h) and GCS-1p::GFP for ROS stress (Fig. 1i). These effects were not due to TIR1 E3 ligase expression, AID-tagging of FASN-1 or auxin treatment as these alone did not enhance the reporters (Extended Data Fig. 3e-g). Because loss of *fasn-1* acted like loss of *ced-3* caspase, we considered the possibility that FASN-1 could be acting downstream of CED-3 in limiting stress responses

### Proteolysis of FASN by caspase attenuates stress response independent of FA synthesis

To confirm that the proteolytic activity of the caspase is required for limiting stress responses, we used a proteolytic dead caspase active-site mutation G360S. This mutant generates full-length CED-3 protein but lacks proteolytic activity. Consistent with *ced-3* null mutants, animals with proteolytic-dead CED-3 also had enhanced stress reporter induction as well as survival under ER, osmotic and ROS stressors (Fig. 2a and Extended Data Fig. 4a-g). This finding suggests that the proteolytic activity of CED-3 is required for regulating stress response. It also explains the role of CED-4 Apaf in stress response as it promotes CED-3 proteolytic activity.

**Fig. 4.**
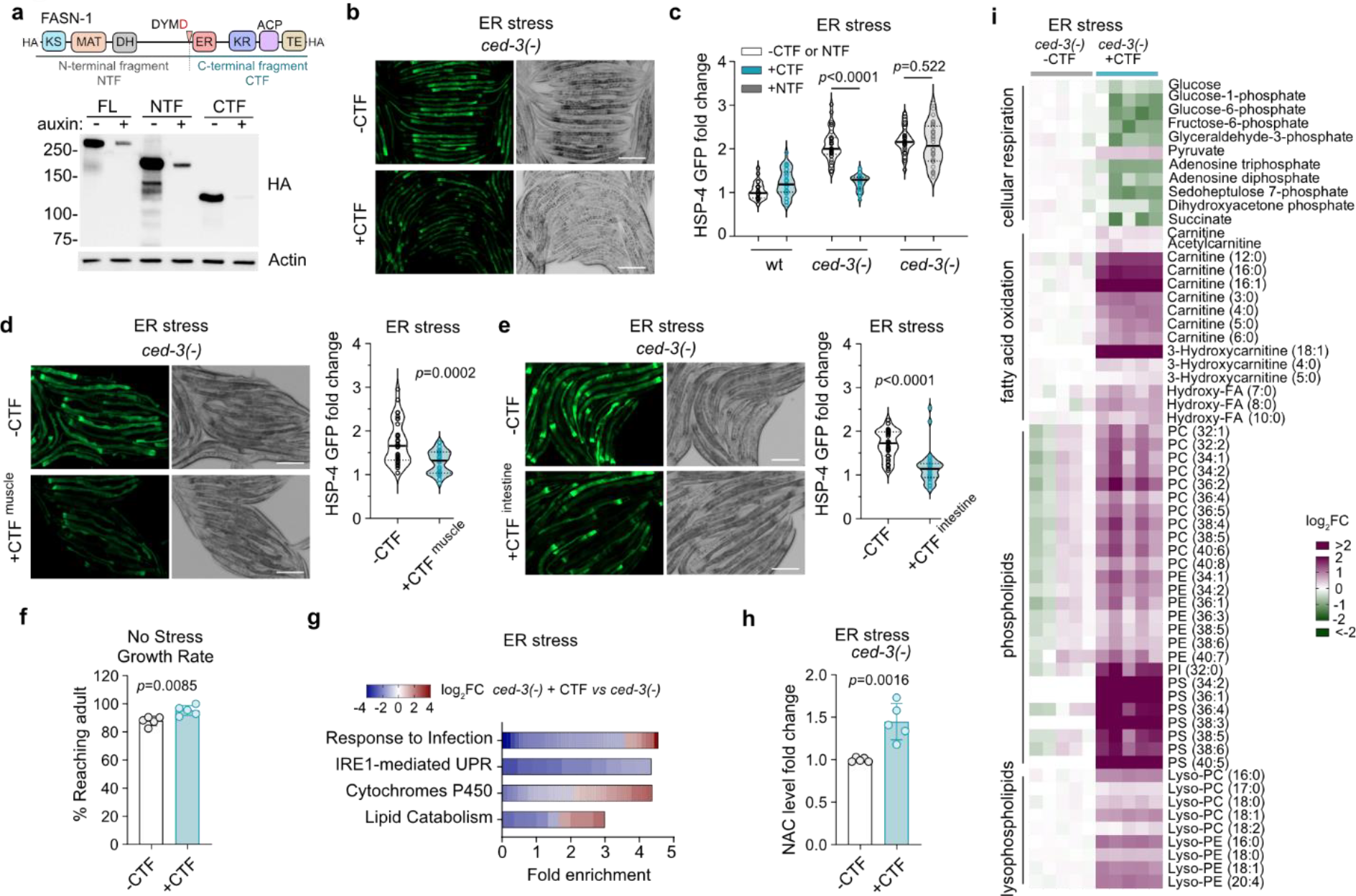
FASN-CTF diminishes chronically-elevated stress-responsive gene expression and metabolic programs. **a**, western blot showing i*n vivo* stabilities of FASN N-terminal fragment (NTF) and C-terminal fragment (CTF). Auxin controlled endogenous FASN-1 (FL), or single copy transgene expressing either NTF or CTF all contain HA tag for detection. **b-c**, HSP-4p::GFP induction (**b**) and quantification (**c**) under ER stress with expression of FASN CTF or NTF in *ced-3(-)*. n=30 animals. **d-e**, HSP-4p::GFP induction under ER stress in *ced-3(-)* animals with expression of FASN CTF in only muscle (**d**) or only in intestine (**e**), n=30 animals. **f**, growth rate of animals with and without expression of FASN-CTF n=5 biological replicates with total of 303-441 animals per condition were assayed,. **g**, FASN-CTF impact on *ced-3*(-) mutant for alterations in gene expression under ER stress. Heatmap shows mRNA Log_2_FC of *ced-3(-)* mutant expressing CTF compared to *ced-3(-)* with no CTF expression. n=3 biological replicates. FDR<0.1 is threshold for enrichment analysis. **h**, N-acetylcysteine (NAC) level in *ced-3(-)* with and without CTF expression. n=5 biological replicates. **i**, impact of FASN-CTF expression on *ced-3(-)* mutant for selected metabolites under ER stress. n=5 biological replicates. Heatmap shows Log_2_FC of *ced-3(-)* mutant expressing CTF compared to *ced-3(-)* with no CTF expression. *p* values were calculated using unpaired ttest (**f,h**) or Mann-Whitney test (**c,d,e**).

To directly test the cleavage of FASN-1 by CED-3, we divided *C. elegans* FASN-1 into 3 large peptide fragments and tested each with *in vitro* cleavage analyses (Fig. 2b). We identified a single CED-3 cleavage site for *C. elegans* FASN-1 in fragment 2 at Asp1593 toward the end of the long non-enzymatic linker region separating the N- and C-termini enzymatic domains (Fig. 2c,d). Like all animal FASN proteins, the human FASN protein has the same domain architecture as *C. elegans* FASN-1 with long intervening non-enzymatic linker. Therefore, we similarly divided *H. sapiens* FASN into 3 large peptide fragments. We found that human Caspase-3 cleaved human FASN fragments 1 and 2 *in vitro* (Fig. 2e). This cleavage leaves an intact C-terminal fragment including most of the ER domain and KR, ACP, and TE domains analogous to the cleavage location for *C. elegans* FASN-1. Furthermore, human Caspase-3 and *C. elegans* CED-3 caspase are known to have similar proteolytic activities. Altogether these results indicate that the cleavage of FASN by caspase may be broadly conserved in metazoans.

To test if the cleavage of FASN-1 by CED-3 is responsible for the enhanced stress responsiveness *in vivo*, we generated a FASN-1 cleavage-resistant mutation D1593E “*fasn-1(D/E*)” in the endogenous locus of *C. elegans fasn-1* using CRISPR mutagenesis. Using Western blot, we first tested FASN-1 expression in wild type, *ced-3(-)* and *fasn-1(D/E*) mutants. Intriguingly, we did not observe any quantitative accumulation of FASN-1 cleavage fragments in *C. elegans* in the absence or presence of stressors (Extended Data Fig. 4h). These results suggest the possibility that only a small fraction of this highly abundant protein is cleaved by CED-3 *in vivo*.

We then tested the impact of *fasn-1(D/E)* on stress response. Similar to the *ced-3(-)* null mutant, the *fasn-1(D/E)* mutant had upregulated HSP-4p::GFP in response to tunicamycin (ER stress, Fig. 2f). This enhanced upregulation was equivalent to *ced-3(-)* (Extended Data Fig. 5a,b) and was confirmed with an independent second *fasn-1(D/E)* CRISPR mutant (Extended Data Fig. 5c). The *fasn-1(D/E)* mutant had enhanced survival comparable to *ced-3(-)* mutants (Extended Data Fig. 5d).

**Fig. 5.**
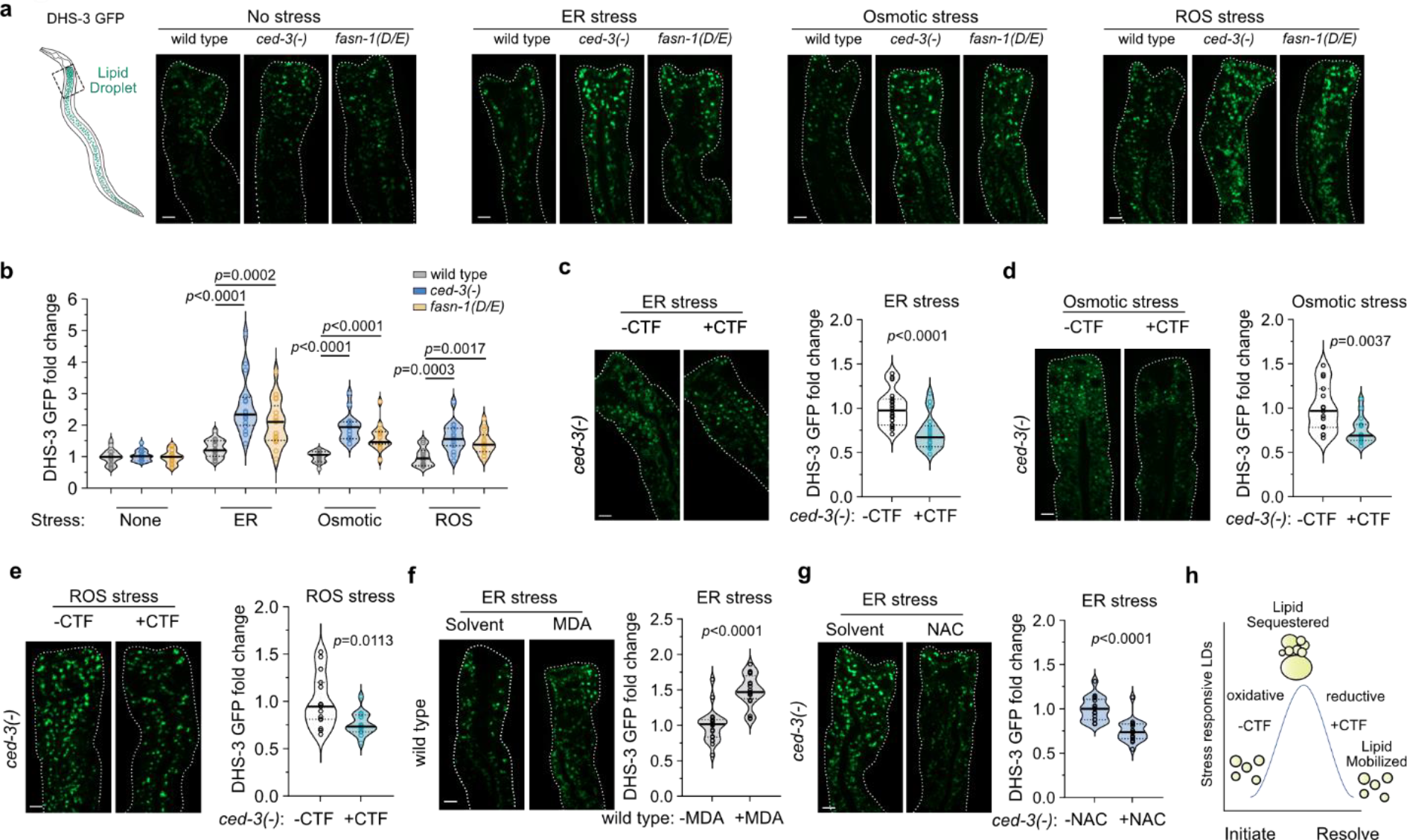
FASN-CTF resolves stress-induced lipid droplets. **a-b**, lipid droplets visualized (**a**) and quantified (**b**) in the same region of intestine by DHS-3::GFP marker intensity under ER, osmotic and ROS stresses. n=15-17 animals per condition. **c-e**, lipid droplets with expression of FASN-CTF in *ced-3(-)* mutant animals under ER (**c**), osmotic (**d**) and ROS (**e**) stresses. n=15-19 animals per condition. **f-g**, lipid droplets with supplementation of malondialdehyde (MDA) in wild type animals (**f**) and N-acetylcysteine (NAC) in ced-3 animals (**g**) under ER stress. n=17-20 animals. **h**, diagram illustrating relationship of lipid droplet accumulation with CTF expression and cellular oxidative status. *p* values were calculated using Mann-Whitney test (**b,c,d,e,f,g**).

We next sought to test whether the *fasn-1(D/E)* mutant would have similar persistent phenotypes with other stressors. Analogous to the heightened ER stress response, the *fasn-1(D/E)* had dramatically enhanced induction of markers for both osmotic stress (NLP29p::GFP) and ROS stress (GCS-1p::GFP) equivalent to *ced-3(-)* when challenged with high salt and paraquat, respectively (osmotic and ROS stresses, Fig. 2f and Extended Data Fig. 5f-j).

Altogether, these data suggest that CED-3 cleaves FASN-1 at the Asp1593 residue to limit pan-stress responsiveness *in vivo*. Because failure of cleavage caused by FASN-1(D/E) mutation phenocopied loss of caspase activity seen with the *ced-3(-)* mutant, we concluded that a low level of caspase cleavage activates a distinct function of FASN to limit pan-stress response.

The saturated fatty acid palmitate (C16) is the major end product of all metazoan FASN enzymes ^6^. To test if *ced-3(-)* and *fasn-1(D/E)* mutations have compromised palmitate synthesis, we performed metabolic labelling with D_2_O to monitor fatty acid synthesis. We found no significant alterations in palmitate synthesis (Fig. 2g) or palmitoleate synthesis (Extended Data Fig. 5k) for *ced-3(-)* or *fasn-1(D/E)* mutants in the with or without ER stress.

To further confirm that the enhanced stress responsiveness of *ced-3(-)* and *fasn-1(D/E)* mutants was not due to a fatty acid deficiency, we supplemented animals with palmitate under ER stress and found that palmitate supplementation did not reduce the heightened HSP-4p::GFP induction in *ced-3(-)* null animals or *fasn-1(D/E)* cleavage-resistant mutants (Fig. 2h). In contrast, extra palmitate enhanced HSP-4p::GFP induction in wild type, *ced-3(-)*, and *fasn-1(D/E)* animals (Fig. 2h). Altogether, these results indicate that the stress resolution function of cleaved FASN is independent of palmitate synthesis.

### FASN cleavage controls redox metabolism

To reveal the metabolic impact of CED-3 caspase on FASN-1, we used untargeted metabolomics in *C. elegans* to analyze the profiles of wild-type, *ced-3(-)*, and *fasn-1(D/E)* strains in normal and stress conditions (Supplementary Dataset 1). To understand the variation for all detected metabolites, we used principal components analysis (PCA). We found that the biggest variance in metabolites (PC1) corresponded to stress treatments including ER (Fig. 3a), osmotic (Fig. 3b) and ROS stressors (Fig. 3c). The second largest variance represented by the PC2 axes corresponded to the different genotypes in the absence of stress. PC2 showed that the *ced-3(-)* and *fasn-1(D/E)* mutants clustered closer to each other (open circles, Fig. 3a-c), suggesting the metabolic profiles of these two mutants are more similar to each other than compared to the wild type. Interestingly, the dispersion amongst the genotypes on PC2 axes collapsed following stress treatments (closed circles, Fig. 3a-c), indicating that all genotypes shared a more similar metabolic profiles under stress. Most of the variability was well-represented in the first 2 principle components for each of the treatments (Extended Data Fig. 6a). The clustering of wild-type with the mutants upon stress treatments suggests a subset of metabolites already altered in *ced-3(-)* and *fasn-1(D/E)* that normally occurs during stress response.

**Fig. 6.**
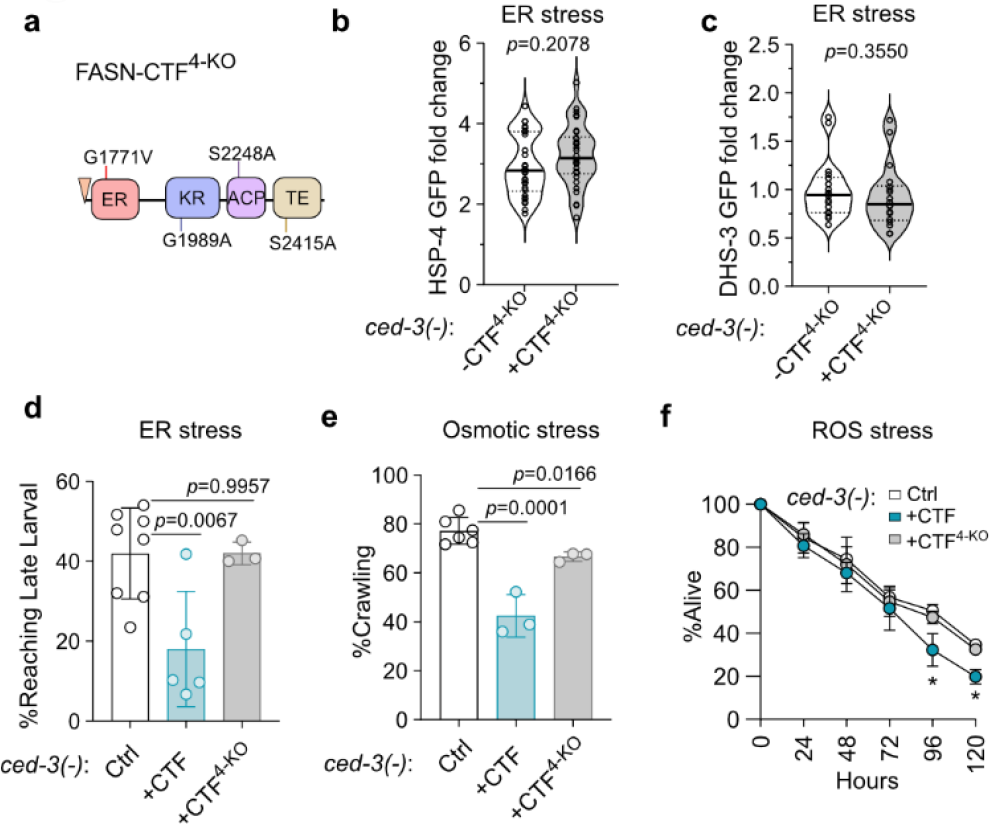
Impact of FASN-CTF enzymatic activity on pan-stress resolution and fitness. **a**, diagram of FASN-CTF with 4 point mutations that inactivate the four conserved enzymatic domains (FASN-CTF^4-KO^). **b**, induction of ER stress marker HSP-4p::GFP with expression of FASN-CTF^4-KO^ under ER stress in *ced-3(-)* mutant animals. n=30 animals. **c**, induction of lipid droplet marker DHS-3p::GFP with expression of FASN-CTF^4-KO^ under ER stress in *ced-3(-)* mutant animals. n=18 animals. **d-f**, artificially driving FASN-CTF expression during stress compromises survival. Fitness cost analyses with either FASN-CTF or FASN-CTF^4-KO^ measured by developmental rate, motility and survival of *ced-3(-)* mutant animals under ER (**d**), osmotic (**e**) and ROS (**f**) stress conditions. n=3-6 biological replicates. Total animals for each bar ranges from 109-298. *p* values, Mann-Whitney test (**b,c**) and unpaired t-test (**d,e,f**).

We further found a strong positive correlation between *ced-3(-)* and *fasn-1(D/E)* mutants compared to wild-type using Pearson’s analysis also supporting that *ced-3(-)* and *fasn-1(D/E)* share similar alterations in metabolites (Extended Data Fig. 6b). Using enrichment analysis of metabolites altered in *ced-3(-)* or *fasn-1(D/E)* mutants, we found a significant enrichment for redox adaptations in the absence of stress for both mutants (Fig. 3d).

We found that most glutathione metabolites detected are depleted in both *ced-3(-)* and *fasn-1(D/E)* mutants in the absence of stress (No Stress, Fig. 3e). Interestingly, the same depletion pattern was observed for wild-type animals under ER and osmotic stress conditions. ROS stress had a somewhat different pattern compared to the other two stressors but still resulted in an overall reduction of the cysteine pool (Fig. 3e). The glutathione pathway modulates cellular redox to mitigate stress responses. Unlike other stressors, ROS stress directly impacts cellular redox and paraquat detoxification directly modulates this pathway.

We also analyzed other enriched pathways for *ced-3(-)* and *fasn-1(D/E)* including carnitine and beta-alanine pathways and found that *ced-3(-)* and *fasn-1(D/E)* shared similar alterations in these pathways but these changes are mostly distinct from stress treatments (Extended Data Fig. 6c,d). Altogether, the metabolomic analyses revealed that both *ced-3(-)* null and the *fasn-1(D/E)* mutants had significant alterations in redox metabolism that resembles a stress-responsive state prior to encounter of a stressor.

### Cellular redox state modulates caspase activity and caspase-dependent stress response

As a Cys protease, caspases can be inhibited by oxidizing species ^17^ and highly oxidizing environments can even suppress caspase-mediated apoptosis ^18, 19^. To assess the relative alteration in caspase cysteine oxidation, we treated primary mouse osteoblasts with hydrogen peroxide overnight. Using the DCP-Bio1 probe that forms adducts with singly-oxidized cysteines (cysteine sulfenic acid), we observed an increase in oxidation of caspase-3 cysteine residues more than 3-fold compared to untreated cells without affecting caspase protein levels (Fig. 3f), suggesting that caspase cysteine residues are sensitive to cellular redox conditions. We next considered if the more oxidized cellular environment under stress treatment would impact caspase activity.

ER stress both results from the accumulation of—and itself generates—reactive oxygen species ^20, 21^. To further assess the direct impact of stress on caspase activity, we tested how ER stress affects CED-3 auto-processing in *C. elegans*. We found that overnight ER stress treatment resulted in diminished CED-3 auto-processing *in vivo* (Fig. 3g). ER stress increased the accumulation of full-length CED-3 protein that corresponded to decreased accumulations of both the p32 and p15 processed subunits (Fig. 3g). This observation indicates that CED-3 has diminished proteolytic capability under ER stress.

For vertebrates, such as mammals, fasting induces oxidative stress in the liver, where FASN expression is enriched. Therefore, we analyzed the effect of fasting on FASN and caspase expression in mouse liver. During the well-fed state, we found FASN processing along with the presence of active caspase-3 whereas both of these were absent during prolonged fasting (Fig. 3h). Altogether, these findings indicate that caspase cysteine oxidation and subsequent inhibition of caspase activity can occur during stress conditions in both cell culture and an animal. The decrease of caspase activity would thereby lead to a heightened stress response similar to the *ced-3(-)* mutation (Fig. 3i).

To further test if modulating cellular redox status directly impacts CED-3 caspase-dependent stress responsiveness, we supplemented *ced-3(-)* animals with antioxidants N-acetylcysteine (NAC, Fig. 3j and Extended Data Fig. 7a,b) or L-carnosine (Fig. 3k and Extended Data Fig. 7c,d) under ER stress conditions. We found that both antioxidant supplements significantly reduced the heightened ER stress response in *ced-3(-)* mutants. These findings suggest that the CED-3 caspase-mediated stress response can be reversed by increasing the cellular reductive capacity.

Conversely, we then considered how organismal stress response would be impacted by shifting the redox state toward an oxidative profile. Lipid peroxidation occurs as a result of oxidative bursts and elevated cellular ROS results in accumulation of lipid-derived reactive aldehydes such as malondialdehyde (MDA). When supplementing animals with MDA, we found that wild type animals had increased response to ER stress similar to *ced-3(-)* mutants without MDA (Fig. 3l and Extended Data Fig. 7e,f). Moreover, MDA did not further increase the elevated response of *ced-3(-)* mutants (Fig. 3m). This result indicates that elevated cellular oxidative reactions augment stress response and loss of CED-3 caspase mimics an elevated oxidative cellular environment. Because *ced-3(-)* null and the cleavage-resistant *fasn-1(D/E)* mutants both had increased amplitude in stress responses, we concluded that CED-3 proteolytic activity may be redox sensitive to modulate the magnitude of response suggesting the possibility of FASN cleavage as a signal to scale down stress response.

### FASN C-terminal Fragment (FASN-CTF) attenuates pan-stress response

To understand the functional outcome of CED-3 cleavage on FASN *in vivo*, we used CRISPR mutagenesis to insert either the N- or the C-terminal cleaved fragment as a single extra copy in *C. elegans* (diagram, Fig. 4a). The N-terminal FASN fragment generated a spectrum of processing bands whereas the C-terminal fragment made one stable product (Fig. 4a). Further analysis of the FASN N-terminal fragment revealed that the linker region at the end of the N-terminal fragment likely contains a degron when exposed by proteolytic cleavage (Extended Data Fig. 8a,b).

To analyze the impact of both FASN N- and C-terminal fragments in stress resolution, we tested the effects of expressing either fragment in *ced-3(-)* null mutants upon stress (Fig. 4b,c). We found that expression of the C-terminal fragment (FASN-CTF) is sufficient to resolve the enhanced stress-responsiveness of the *ced-3(-)* mutant while the N-terminal fragment is not (Fig. 4b,c). Moreover, tissue-specific expression of FASN-CTF in either the intestine or muscle (Extended Data Fig. 8c) also decreased the heightened induction of the intestinal HSP-4p::GFP of the *ced-3(-)* mutant (Fig. 4d,e and Extended Data Fig. 8d-g). Therefore, we concluded that FASN-CTF can function cell non-autonomously to decrease stress response.

Because FASN is an essential gene and loss of FASN-1 causes developmental arrest in *C elegans* ^12, 22^, we wanted to determine if the FASN-CTF works to inhibit intact FASN function. To assess fitness cost, we measured the growth rate of animals expressing FASN-CTF and found no growth delay (Fig. 4f), suggesting FASN-CTF functions to decrease stress response without affecting the essential *de novo* fatty acid synthesis function of FASN during development.

### FASN-CTF promotes anti-inflammatory profile

To further investigate the impact of FASN-CTF on gene expression, we performed mRNA-Seq analysis (Supplementary Dataset 2). We found that with ER stress, expressing the CTF in *ced-3(-)* decreased expression of genes enriched for pathogen defense and IRE1-mediated UPR (Fig. 4g and Extended Data Fig. 8h,i). We also saw mixed alterations in genes enriched for Cytochrome P450 and lipid metabolism. Genes in pathogen defense and UPR were almost all down-regulated by FASN-CTF (Fig. 5g and Extended Data Fig. 8h,i), suggesting a strong anti-inflammatory-type response.

We also analyzed the metabolic impact of expressing FASN-CTF on *ced-3(-)* under ER stress using targeted metabolomics (Supplementary Dataset 3). Interestingly, we found elevated N-acetylcysteine anti-oxidant with FASN-CTF expression (Fig. 4h). This finding was consistent with our observation that supplementing N-acetylcysteine reduces the heightened ER stress response in *ced-3(-)* null mutants (Fig. 3j).

We also found that *ced-3(-)* null mutants with ectopic expression of FASN-CTF had altered enrichments for metabolites in multiple energy production pathways (Extended Data Fig. 8j-k). Specifically, when expressing FASN-CTF, most metabolites tested decreased in glycolysis and TCA cycle pathways (Fig. 4i). Conversely, almost all carnitine species, hydroxy-fatty acids and phospholipids tested were increased with FASN-CTF expression (Fig. 4i), suggesting enhanced fatty acid oxidation.

Altogether, we found that expressing FASN-CTF attenuates stress-responsive gene programs, elevates reducing metabolites and importantly, shifts energy metabolism from cellular respiration to fatty acid oxidation with elevated soluble lipids. Therefore, we considered the possibility that FASN-CTF impacts lipid mobilization.

### FASN-CTF resolves stress-induced lipid droplets

Lipid droplets not only serve as readouts of stress responses but also play essential roles in mitigating stress responses, integrating lipid metabolism with energy homeostasis, supporting proteostasis during stress responses and sequestering reactive oxidizing species ^23-25^. To observe intestine-specific lipid droplets, we used the DHS-3::GFP fusion protein as a previously established marker in *C. elegans* ^26^. Without stress treatment, *ced-3(-)* null and *fasn-1(D/E)* mutants had no effect on basal lipid droplet intensity as visualized by DHS-3::GFP (No Stress, Fig. 5a). With challenges from ER, osmotic and ROS stress, *ced-3(-)* and *fasn-1(D/E)* mutants had a marked increase in lipid droplet intensity (Fig. 5a,b) but had no impact on the DHS-3::GFP protein levels (Extended Data Fig. 9a), suggesting increased accumulation of lipid droplets in *ced-3(-)* and *fasn-1(D/E)* mutants with stress treatments.

We next tested whether expressing FASN-CTF would impact lipid droplet dynamics of *ced-3(-)* mutants. We found that the FASN-CTF was able to ameliorate the heightened lipid droplet accumulation in *ced-3(-)* mutants with ER (Fig. 5c), osmotic (Fig. 5d), and ROS stressors (Fig. 5e). When expressing FASN-CTF in wild type animals, there were no differences in lipid droplets with ER, osmotic, or ROS stressors (Extended Data Fig. 9b-d). This was expected given that wild-type animals have functional CED-3 and FASN-1.

Given the ability of the FASN-CTF to promote cellular reductive capacity, we further tested how lipid droplet dynamics were affected by altering cellular redox conditions. We found that supplementing wild type animals with the lipid aldehyde metabolite MDA enhances lipid droplet accumulation under ER stress (Fig. 5f) whereas supplementation with the anti-oxidant NAC ameliorated the lipid droplets accumulation in *ced-3(-)* mutants (Fig. 5g). These findings indicate that expressing FASN-CTF had similar effects as restoring cellular redox on lipid droplet dynamics (Fig. 5h).

### FASN-CTF catalytic activity signals stress resolution

Previous work identified point mutations that disrupt mammalian FASN enzymatic activity ^27^. We introduced a transgene of FASN-CTF with point mutations of the highly conserved residues to inactivate each of the 4 domains in the FASN-CTF and referred to as FASN-CTF^4-KO^ (Fig. 6a). These residues were identified by homology to the human FASN (Extended Data Fig. 10a). The FASN-CTF and FASN-CTF^4-KO^ were expressed at similar levels and were both responsive to auxin-induced degradation (Extended Data Fig. 10b). Analogous to FASN-CTF, expression of FASN-CTF^4KO^ did not delay development under normal conditions suggesting it also does not impact endogenous FASN-1 function (Extended Data Fig. 10c). In contrast to FASN-CTF, expression of the FASN-CTF^4-KO^ did not suppress the ER stress response as visualized by continued expression of HSP-4p::GFP (Fig. 6b and Extended Data Fig. 10d). Additionally, FASN-CTF^4KO^ also did not ameliorate heightened lipid droplet accumulation (Fig. 6c and Extended Data Fig. 10e). These results suggest that catalytic activity of FASN-CTF is required for its function in attenuation of stress response.

To distinguish whether the FASN-CTF functions more as an effector to mitigate stress or more as a signal to scale down stress responses, we assayed developmental progression under ER stress. We found that expression of FASN-CTF led to more stalled development for the *ced-3(-)* mutants in the presence of ER stress (+CTF, Fig. 6d). In contrast, the FASN-CTF^4-KO^ behaved similar to *ced-3(-)* mutants (+CTF^4-KO^, Fig. 6d). Similarly, expression of FASN-CTF led to less motility under osmotic stress (Fig. 6e) and less survivors under ROS stress (Fig. 6f). Because expressing FASN-CTF decreased overall fitness of *ced-3(-)* mutants under diverse stress conditions, we concluded that the presence of FASN-CTF prior to stress mitigation compromises survival as it provides a premature stress resolution signal.

In conclusion, we propose that FASN-CTF is generated by caspase cleavage in a redox-dependent manner. Under stress conditions, more oxidative cellular environment renders CED-3 caspase less active, leading to increased stress response. Once stress is mitigated, less oxidative cellular environment re-activates caspase and generates FASN-CTF through proteolytic cleavage of FASN. FASN-CTF functions as an *all clear* signal to initiate a stress response attenuation program. As an *all clear* signal, FASN-CTF is sufficient to promote lipid mobilization and fatty acid oxidation as well as down regulate expression of both UPR and innate immunity stress response genes. As a result, physiological adaptations are re-programmed for a non-stress state to support growth and homeostasis (Extended Data Fig. 10f).

## Discussion

Lipid biosynthesis itself has an intriguing function in activating stress responses. Compromised *de novo* fatty acid synthesis by mutation of *fasn-1* enhanced the expression of anti-microbial genes in *C. elegans* ^22^ and oxidative stress response genes in yeast ^28^. However, lipid biosynthesis by FASN-1 was shown to be required to initiate the mitochondria-to-cytosolic stress response triggering HSF-1 and DVE-1 transcriptional programs ^29^. Moreover, FASN is required for diet-induced inflammatory signaling ^30^ as well as axonal regeneration following nerve injury ^31^. These seemingly contradictory findings suggest a complex role for FASN in stress responses which could be explained by roles in both lipid synthesis as well as signaling that we find.

Alterations in lipid accumulation activate SKN-1 Nrf which itself is critical in lipid homeostasis ^32^. Redox-sensitive transcriptional programs are essential to initiate an anti-oxidant response that mitigates oxidative challenges ^33, 34^. Thus, a reversible sensor acting at the post-translational level would reinforce dynamic sensing of stress responses allowing for rapid sequestration of energy stores. Limiting lipid peroxidation is of paramount importance for cell survival during stress ^7^. Thus, it is intriguing that the fatty acid-synthesizing enzyme FASN would also have a stress-sensing function, signaling the all clear after stressful encounters have been attenuated. Additionally, a shift in redox is a general feature of many stressful and pathogenic states making redox restoration coupled with lipid droplet dynamics an effective adaptive response for stress mitigation. Moreover, accurately gauging a response proportional to the challenge without mounting an excessive response allows cellular energy supplies to be conserved. Redox signaling is complex in that the redox factors can take on multiple roles ^35, 36^.

Efforts to map the cysteinome have revealed an array of oxidized cysteine species and complex roles of cysteine redox in signaling, gene expression, cellular growth and stress responses ^37, 38^. Depending on the oxidation state, multiple cysteine oxidation species are reversible ^39, 40^. However, some cysteine oxidation forms are irreversible ^39, 40^. Following redox resolution by mechanisms including SKN-1 Nrf transcriptional regulation ^33, 34, 41^, NADPH oxidase signaling ^42^ and subsequent sulfide production ^43^, cysteine redox balance is collectively restored and the caspase active-site cysteine would be reactivated and allow for normal lipid droplet dynamics. Therefore, caspase makes for an intriguing sensor of oxidation state where mitigation of reactive oxidative species could restore activity and allow for resumed proteolytic activity.

Animals have evolved signaling mechanisms ranging from lipid-derived ligands to phosphorylation cascades to protein-protein interactions ^44-50^. In many cases, the signaling molecule or interaction responsible for these functions are present at minute levels, yet have potent roles as pro-development or stress-activating cues. Solving each of these elegant signaling mechanisms as multi-step cascades represented considerable effort with numerous studies. Our current findings reveal an unexpected role of FASN in detecting stress levels via a redox-sensitive, caspase-dependent proteolytic generation of the FASN-CTF. We find that generation of the FASN-CTF is a strong stress-resolution cue. Generation of the FASN-CTF can even be detrimental with forced expression of FASN-CTF without mitigation of the stressor. The duality of FASN as a biosynthetic enzyme and a generator of the FASN-CTF stress-resolution cue suggests FASN may have a broader role as a signaling hub integrating basic metabolic demands and stress inputs.

## Acknowledgements

We thank David Mangelsdorf, Steven Kliewer, Duojia Pan, Melanie Cobb, Joel Goodman, Elliott Ross, Matthew Sieber, Mike Henne, Michael Reese, James Collins, and all members of the Weaver lab for helpful discussions; the CGC (funded by NIH Office of Research Infrastructure Programs [P40 OD010440]) for materials; WormBase, MetaboAnalyst, Small Molecule Pathway Database (SMPDB) and UniProt databases. Targeted metabolomics screen was provided by the CRI Metabolomics Core. mRNA-seq was performed by the McDermott Center Sequencing Core and data analysis was provided by the McDermott Center Bioinformatics Lab. This work is supported by Welch Foundation grant I-2022-20190330 (BPW), National Institutes of Health grants R35GM133755 (BPW), R35CA22044901 (RJD), R01AR076325 (CMK), R01AR071967 (CMK), and Investigator funding through HHMI (RD). The funders had no role in study design, data acquisition, decision to publish, or preparation of the manuscript.

## Author contributions

Conceptualization, HW, YMW, and BPW; Methodology and Investigation, HW, YMW, CY, YZ, GH, CMK, and BPW; Formal Analysis, HW, YMW, CY, YZ, GH, CMK, RJD and BPW; Writing, HW, YMW, RJD, and BPW; Supervision of Study, YMW, RJD, and BPW; Funding Acquisition, RJD, CMK, and BPW.

## Declarations of interests

### Vertebrate animals

All animal use was approved by the Institutional Animal Care and Use Committee (IACUC) at the University of Texas Southwestern Medical Center at Dallas, Texas.

## Data and software availability

All mRNA-Seq data are deposited in GEO accession GSE226048

## Supplementary Information

- **Extended Data Figures with legends**
- **Methods**
- **Supplementary Dataset 1, Untargeted metabolomics**
- **Supplementary Dataset 2, mRNA seq of FASN-CTF vs ced-3**
- **Supplementary Dataset 3, Targeted metabolomics of FASN-CTF vs ced-3**

